# Real-time Noise-suppressed Wide-Dynamic-Range Compression in Ultrahigh-Resolution Neuronal Imaging

**DOI:** 10.1101/2021.09.29.462090

**Authors:** Bhaskar Jyoti Borah, Chi-Kuang Sun

## Abstract

With a limited dynamic range of an imaging system, there are always regions with signal intensities comparable to the noise level, if the signal intensity distribution is close to or even wider than the available dynamic range. Optical brain/neuronal imaging is such a case where weak-intensity ultrafine structures, such as, nerve fibers, dendrites and dendritic spines, often coexist with ultrabright structures, such as, somas. A high fluorescence-protein concentration makes the soma order-of-magnitude brighter than the adjacent ultrafine structures resulting in an ultra-wide dynamic range. A straightforward enhancement of the weak-intensity structures often leads to saturation of the brighter ones, and might further result in amplification of high-frequency background noises. An adaptive illumination strategy to real-time-compress the dynamic range demands a dedicated hardware to operate and owing to electronic limitations, might encounter a poor effective bandwidth especially when each digitized pixel is required to be illumination optimized. Furthermore, such a method is often not immune to noise-amplification while locally enhancing a weak-intensity structure. We report a dedicated-hardware-free method for rapid noise-suppressed wide-dynamic-range compression so as to enhance visibility of such weak-intensity structures in terms of both contrast-ratio and signal-to-noise ratio while minimizing saturation of the brightest ones. With large-FOV aliasing-free two-photon fluorescence neuronal imaging, we validate its effectiveness by retrieving weak-intensity ultrafine structures amidst a strong noisy background. With compute-unified-device-architecture (CUDA)-acceleration, a time-complexity of <3 ms for a 1000×1000-sized 16-bit data-set is secured, enabling a real-time applicability of the same.

## Introduction

Optical microscopy [1-3], a widely used technique for neuronal imaging, has been helping researchers over the past several decades visualize and understand various neurological disorders, brain-functions and - dysfunctions. Neuronal structures [4-6] often show a wide variation in structural texture and signal strength. For instance, while imaging a neuron, the soma, i.e., the cell body, is often expected to be order-of-magnitude brighter than the adjacent fiber structures, such as axons and dendrites, which can be thinner than even a micron in diameter. This essentially leads to a broad enough signal intensity distribution, which is less likely to be properly digitized and visualized with a limited dynamic range of a state-of-the-art acquisition and display system. Additionally, various optical, electrical, and other environmental factors often result in a noisy background which contaminates the weaker intensity signals and worsens the signal-to-noise ratio (SNR) as well as the contrast ratio. Aside from this, to prevent a possible photobleaching and/or phototoxicity [7], i.e., a laser-induced damage to a tissue under observation, it is often recommended to maintain a low enough excitation power, which in turn further deteriorates the signal strengths from the ultrafine neuronal structures. Likewise, while performing a three-dimensional (3D) optical sectioning of a deep volumetric tissue sample [8-9], due to frequency dependent scattering and absorption issues [10-11], the signal-of-interest tends to degrade even further as one penetrates deeper into the tissue. The issue gets even worse due to non-ideal optical performance of a scanning system. Particularly for a large imaging area in a mesoscopic imaging system [12-15], the optical aberrations [16] become prominent towards the edges and corners, unavoidably leading to non-uniform excitation and detection efficiencies across the FOV. This essentially further reduces the signal strengths of the weaker structures residing at the off-axis locations. As a matter of fact, the weaker intensity signals from the ultrafine neuronal structures tend to get closer and closer to the noisy background, even when the bright pixels of the images are almost saturated. It thus becomes a challenging task to retrieve such weaker structures with an adequate SNR and a high contrast-ratio together with the brighter ones, all amidst a strong noisy background. A straightforward attempt to enhance the contrast of such weak structures might lead to amplification of noise, and additionally, it is very likely that the brightest structures tend to saturate due to a limited dynamic range of the system.

Several hardware-based techniques have been reported over the years to address the dynamic range limitation in optical microscopy, which can locally enhance the weaker structures while preventing brighter structures from saturating. One promising solution is to regulate the excitation power in real-time. In such an approach, a feedback mechanism is utilized which monitors the emerging signal strength and accordingly provides a suitable feedback to a tunable excitation source to regulate the excitation power [17-21]. Another approach is to employ a real-time high-dynamic-range (HDR) imaging [22-24], which collects multiple low-dynamic-range (LDR) images over multiple optically-separated detection channels, and subsequently fuses them together to form the HDR image. However, implementing these techniques requires dedicated hardware configurations. For instance, a proper feedback-electronic-circuit, a tunable excitation source, and at least one dedicated channel for monitoring the output signal strength are required for a regulated/ controlled/ adaptive illumination to work. A typically slower response due to electronic limitations might however lead to a poor effective bandwidth especially when each digitized pixel is required to be illumination optimized. A lower effective bandwidth might in turn result in *aliasing* [25-26], i.e., an irreversible loss of digital resolution, especially when concerning a high spatial resolution over a large millimeter-scale FOV. Besides, such a method is often not immune to noise-amplification while locally enhancing a weak-intensity structure. Likewise, in case of real-time HDR imaging, multiple channels are dedicated to detect the same spectral regime with different signal strengths, and thus multi-spectrum detection for multi-color imaging becomes complex in the context of optical design implementation.

Quite a few software-based approaches have been developed either to enhance the contrast and/or sharpness of an image while minimizing the non-uniform-illumination issue, or to perform various image-segmentation operations. A few of the techniques employ subtraction of a mask/layer, which involves analyzing the relevant image or image-stack, predicting a subtraction mask(s) accordingly, and finally subtracting the same from the original image. Traditional and modified unsharp masking methods [27-32], no neighbor/ nearest neighbor method [33], rolling ball/ sliding paraboloid background subtraction [34-35] are some of the popular techniques in this regard. Recently, we have reported a modified unsharp masking algorithm [36], which was dedicated to suppress high-frequency noises in a background while mostly preserving useful information. This approach was however limited to suppression of noise, and was further constrained by choices of multiple controlling parameters. There are several other noteworthy algorithms/ techniques [37-49] which have been improving the image quality over the past several years. Aside from them, another widely used approach to eliminate high-frequency noises which can help to improve both SNR as well as contrast-ratio is to perform certain morphological operations [50], such as, erosion and opening. However, while performing an erosion operation, it is quite possible that certain high-frequency useful information from the image gets removed. A subsequent dilation operation (i.e., an opening operation involving an erosion followed by a dilation) might no longer be able to regenerate the lost information, and thus might lead to an irreversible resolution loss. Another promising approach to enhance the image contrast is to perform a traditional, or an adaptive histogram equalization (HE, or AHE) [51-56]. However, when an image consists of a noisy background, HE, or AHE might lead to an amplification of noise and thus SNR might degrade significantly. An improved version of AHE, i.e., a contrast limited adaptive histogram equalization (CLAHE) [57-59] is a widely used state-of-the-art local contrast enhancement technique which can limit the noise-amplification issue significantly. However, the issue of noise-amplification might still persist in case of an optical microscopy image due to the fact that a significant portion of the image might not possess useful information, but might consist of a strong noisy-background only, which we do not intend to amplify or enhance.

In this paper, we report a dedicated-hardware-free method to achieve a real-time noise-suppressed wide-dynamic-range compression and/or contrast enhancement to virtually mimic the effect of hardware-based adaptive illumination technique. To demonstrate our idea, we performed large-FOV Nyquist-satisfied (aliasing-free) wide-bandwidth two-photon fluorescence imaging of brain/neuronal structures at multiple excitation wavelengths with our custom developed multiphoton optical microscopy (MPM) system [60]. The effectiveness of our method is validated by retrieving weak-intensity ultrafine neuronal structures amidst a strong noisy background, while achieving simultaneous improvements to the signal-to-noise ratio (SNR), signal-to-background ratio (SBR), and contrast-ratio. To secure a real-time applicability, we implement the method via Graphics Processing Unit (GPU)-assisted NVIDIA’s Compute Unified Device Architecture (CUDA)-acceleration with a <3 ms of time-complexity for a typical 1000×1000-sized 16-bit input data-set.

## Results

### Implementation and mathematical formulation/ description of the proposed method

A basic block diagram representation of our data acquisition, processing, and display strategies is presented in Figure 1. A simple laser-scanning fluorescence detection unit is shown in the red-dashed box in Figure 1(a), where EXC and DBS respectively stands for a raster-scanning excitation beam emerging from a laser source which gets focused onto a biological sample by means of an objective lens, and a dichroic beam splitter for separating and guiding the generated fluorescence signal as indicated by the green arrows towards an electronic detection unit comprising of a photomultiplier tube (PMT), a transimpedance amplifier, and a digitizer with a high enough resolution and an adequate bandwidth. Once a frame is scanned and data becomes ready, the raster-scanning system is free to acquire the next frame, provided the number of pending frame(s) to process is not more than 1 and adequate data buffers are available. Please note that most of the state-of-the-art digitizers are capable of providing high-bit-depth data, and therefore we will demonstrate our approach for 16-bit data sets. Up-to this point, to maintain the data-acquisition process, single or multiple CPU-threads can be dedicated as per choice of the digitizer and relevant Application Programming Interfaces (APIs) available.

**Figure 1.**
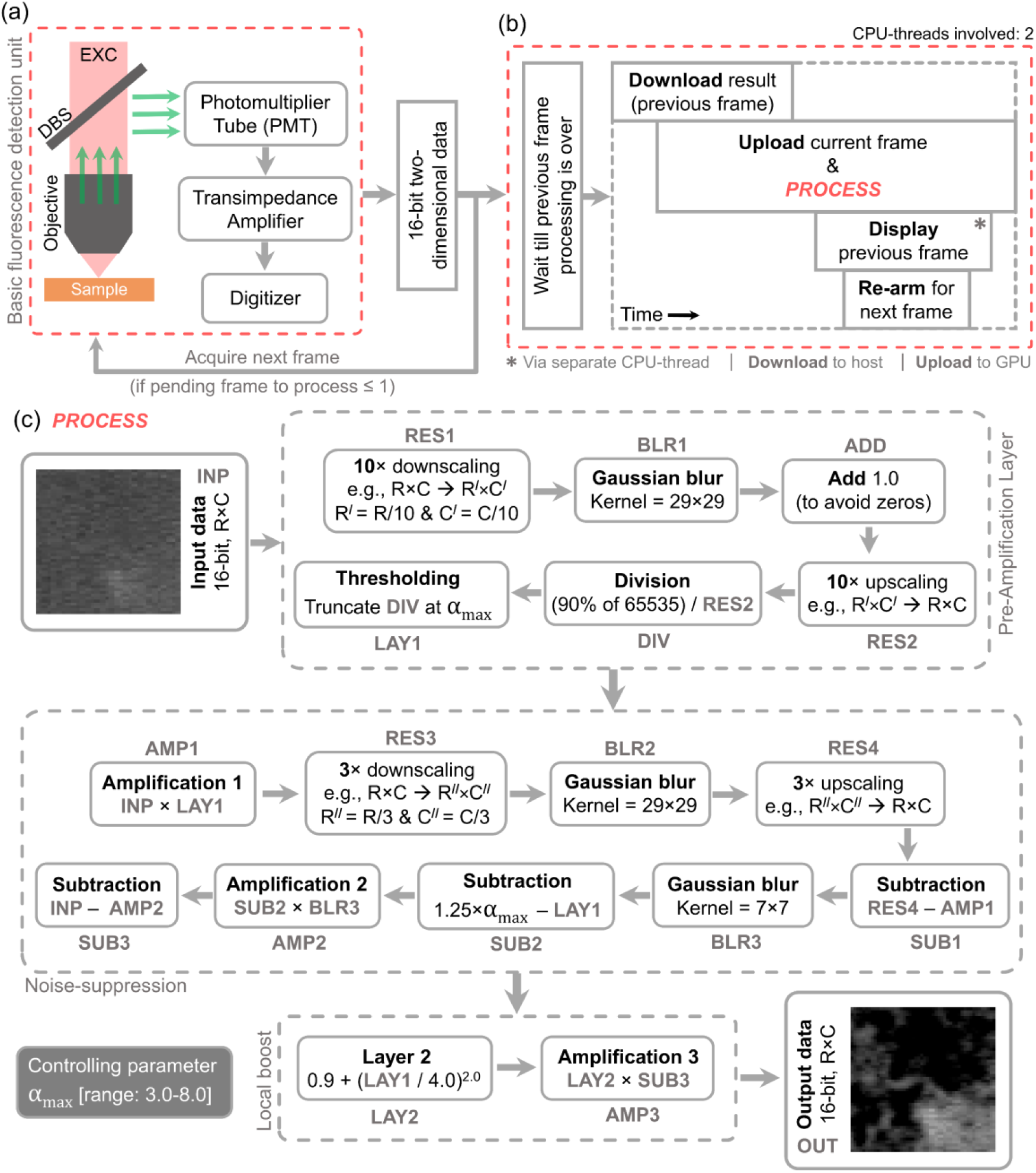
Block diagram representation of data acquisition, process and display strategies. (a) A fluorescence detection unit. EXC: raster-scanning excitation beam. DBS: dichroic beam splitter. EXC gets focused onto sample via objective lens. DBS separates and guides fluorescence signal (green arrows) towards detection unit comprising photomultiplier tube (PMT), transimpedance amplifier, and digitizer. (b) To ensure no pending process, the digitizer asynchronously downloads previous frame, asynchronously uploads current frame, schedules PROCESS-tasks for current frame, displays previous frame via a separate thread, and re-arms. (c) Process tasks. INP: 16-bit input data, RES1: 10× downscaled INP, BLR1: 29×29-kernel gaussian-blurred RES1, ADD: BLR1 added with 1.0, RES2: upscaled ADD, DIV: (90% of (216-1)) divided by RES2, LAY1: truncated DIV at αmax, AMP1: INP multiplied with LAY1, RES3: 3× downscaled AMP1, BLR2: 29×29-kernel gaussian-blurred RES3, RES4: upscaled RES3, SUB1: subtraction of AMP1 from RES4, BLR3: 7×7-kernel gaussian-blurred SUB1, SUB2: subtraction of LAY1 from 1.25 times αmax, AMP2: SUB2 multiplied with BLR3, SUB3: subtraction of AMP2 from INP, LAY2: square of one fourth of LAY1 added with 0.9, AMP3: LAY2 multiplied with SUB3; controlling parameter recommended range: 3.0 ≤ αmax ≤ 8.0.

To comprehend the implementation steps, please refer to Figure 1(b). Once a 16-bit two-dimensional (2D) data set is acquired, we first ensure that previous frame has been processed/displayed, and then start downloading the previous processed data from GPU in an asynchronous manner. Immediately, the newly acquired frame is asynchronously uploaded to the GPU and the whole set of subsequent *PROCESS*-tasks are scheduled thereafter. Following this step, we ensure the scheduled download is complete and display the downloaded result via a different CPU-thread. In the meantime, the main thread re-arms for the next frame. Figure 1(c) illustrates the *PROCESS*-tasks in terms of a simplified block diagram. For a mathematical formulation, let us first assume a noise-affected wide-dynamic-range (low-contrast) 16-bit image *f*(*r, c*) marked as INP in Figure 1(c) with R×C pixels, where *r* and *c* stand for row and column positions, respectively. Applying a 10× downscaling to *f*(*r, c*) we obtain *f*^*D*^ (*r*^/^, *c*^/^) with a reduced pixel number of R^/^×C^/^, as depicted in Equation (1) and RES1 in Figure 1(c). Note that for all R^/^×C^/^ -sized images, *r*^/^ and *c*^/^ stand for row and column positions, respectively. Now, a 29×29 -kernel gaussian blurring is applied to *f*^*D*^ (*r*^/^, *c*^/^), and the blurred result is marked as BLR1 in Figure 1(c). To avoid division-by-zero in the next step, each pixel-value of BLR1 is added with 1.0, and the resultant image is denoted as *l*(*r*^/^, *c*^/^) in Equation (2) and ADD in Figure 1(c). With a bilinear interpolation, *l*(*r*^/^, *c*^/^) is resized back to R×C pixels and the interpolated result is denoted as *l*^*U*^ (*r, c*) in Equation (3) and RES2 in Figure 1(c). An inverse of each *l*^*U*^ (*r, c*) -pixel-value is now multiplied with 90% of the maximum allowed intensity, i.e., 0.9×(2^16^ −1) for a 16-bit image, and the result is given as *d*(*r, c*) in Equation (4). Note that *d*(*r, c*) involves nothing but a division operation, and is marked as DIV in Figure 1(c). Each pixel value of *d*(*r, c*) above *α*_*max*_ is truncated to *α*_*max*_, and the resultant layer is denoted as *α*(*r, c*) in Equation (5) and LAY1 in Figure 1(c).

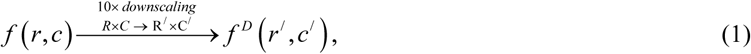

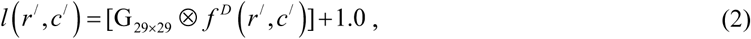

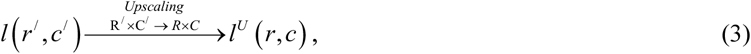

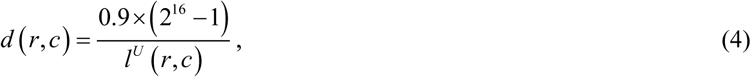

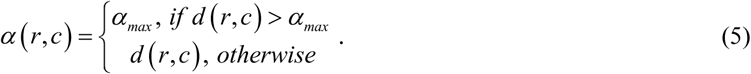

Now, a pixel-to-pixel multiplication of *f*(*r, c*) and *α*(*r, c*) is performed, and thereby AMP1 in Figure 1(c) and *g*(*r, c*) in Equation (6) are obtained. With a 3×3 pixel-binning on *g*(*r, c*), we obtain RES3 in Figure 1(c) and *g*^*D*^ (*r*^//^, *c*^//^) in Equation (7) with a reduced pixel number of R^//^×C^//^, where *r*^//^ and *c*^//^ stand for row and column positions, respectively. A 29×29 -kernel gaussian blurring is applied to *g*^*D*^ (*r*^//^, *c*^//^), and the blurred result is obtained as L(*r*^//^, *c*^//^) in Equation (8) and BLR2 in Figure 1(c). With a bilinear interpolation, L(*r*^//^, *c*^//^) is resized back to R×C pixels and the interpolated result (RES4 in Figure 1(c)) is denoted as L^*U*^ (*r, c*) in Equation (9). Now, a subtraction of L^*U*^ (*r, c*) and *g*(*r, c*) is performed whose result is marked as SUB1 in Figure 1(c), and a 7×7 -kernel gaussian blurring is subsequently applied to SUB1. The blurred result thus obtained is depicted as L^/^ (*r, c*) in Equation (10) and BLR3 in Figure 1(c).

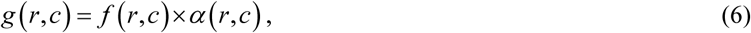

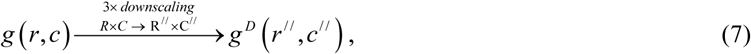

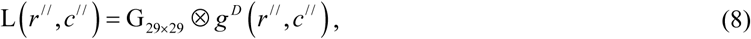

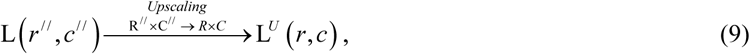

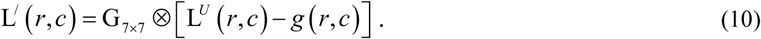

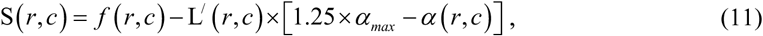

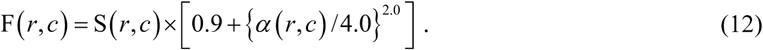

Based on *α*(*r, c*), a modified layer is obtained as [1.25×*α*_*max*_ − *α*(*r, c*)] which is marked as SUB2 in Figure 1(c). Each pixel-value of L^/^ (*r, c*) is now multiplied with each corresponding pixel-value in this layer SUB2, and the result is marked as AMP2 in Figure 1(c). Subsequently, AMP2 is subtracted from *f*(*r, c*) to obtain a noise-suppressed output S(*r, c*), as depicted in Equation (11) and SUB3 in Figure 1(c). Based on the same *α*(*r, c*), another layer LAY2 in Figure 1(c) is now obtained as [0.9 +{*α*(*r, c*) / 4.0}^2.0^], and finally each pixel-value of S(*r, c*) is multiplied with each corresponding pixel-value of LAY2, and thereby the noise-suppressed contrast-enhanced output is obtained in 16-bit format (and optional 8-bit format for display purpose), denoted as F(*r, c*) in Equation (12) and OUT in Figure 1(c).

### Demonstration of the method via two-photon fluorescence images of neuronal structures

To demonstrate our approach, we acquire two-photon fluorescence images of a Nav1.8-tdTomato-positive mouse dorsal-root-ganglion (DRG) section comprising soma and fine axon fibers, and a coronal section from a Thy1-GFP-positive mouse brain cortex region comprising axons, dendrites, and dendritic-spines, excited at central wavelengths of 1070 nm and 919 nm (70 MHz, <60 fs), respectively, with an average excitation power of <40 mW in each case (refer to *STAR Methods*). Figures 2(a) and (c) depict two two-photon images of the Nav1.8-tdTomato and Thy1-GFP samples, respectively, each with an FOV of 1×1 mm^2^, scale bar of 150 µm, however each with a poor SNR, SBR, and contrast-ratio. To improve the same, the proposed method is applied and based on visual response, the value of *α*_*max*_ is adjusted to (a) 8.0 and (c) 6.0, and the corresponding noise-suppressed contrast-enhanced results are depicted in Figures 2(b) and (d), respectively. Please note that we maintain pixel-sizes of of 182 nm and 167 nm for the excitation wavelengths of 1070 nm and 919 nm, respectively. We thus satisfy the Nyquist-Shannon criterion [61-62] and ensure an aliasing-free imaging [25-26] for our 0.95 numerical-aperture (NA) objective lens with diffraction limited two-photon spatial resolutions of 429 nm and 368 nm for the respective excitation wavelengths.

**Figure 2.**
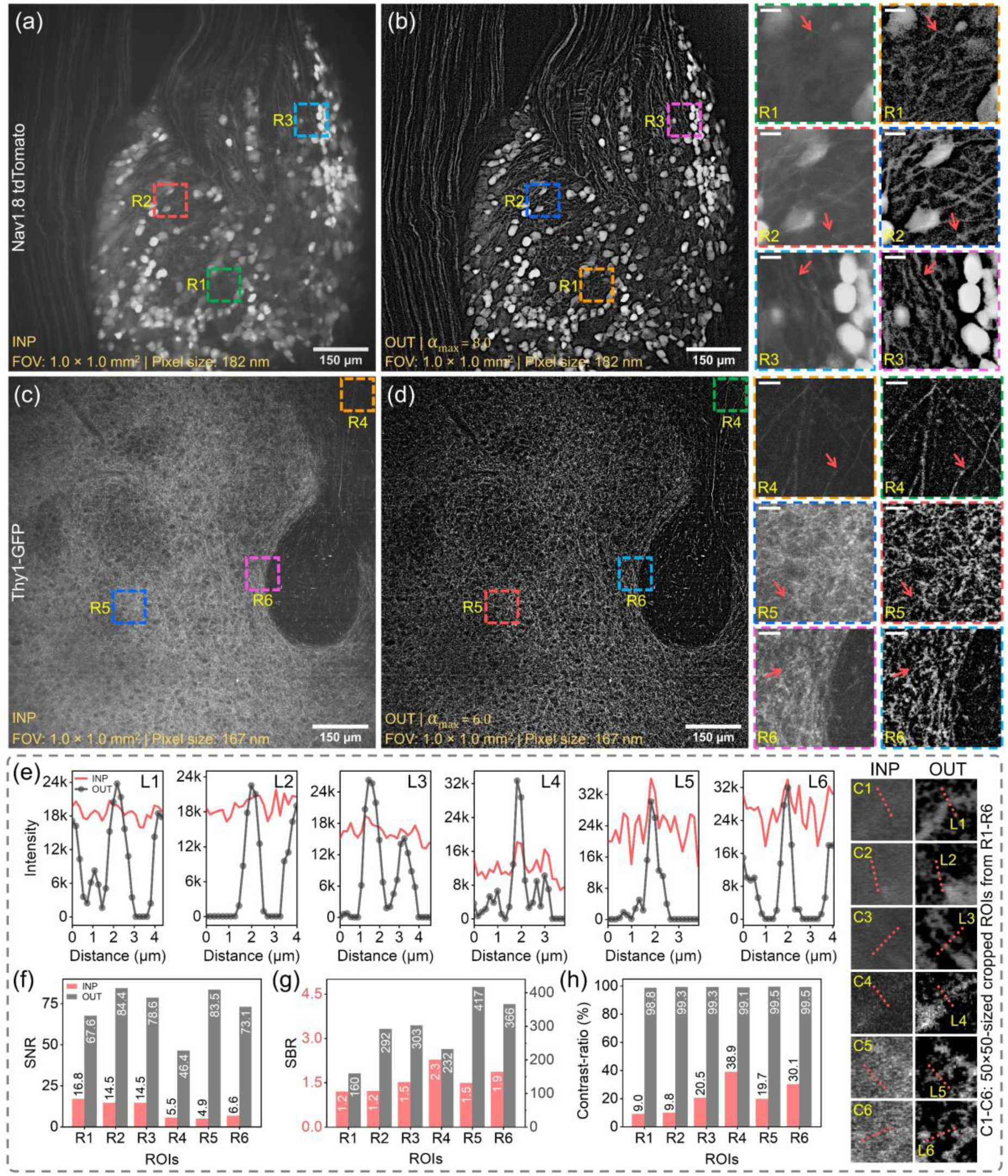
Demonstration of the proposed noise-suppressed wide-dynamic-range compression method in fluorescence microscopy imaging. (a) & (c) Two-photon fluorescence images of Nav1.8-tdTomato-positive mouse dorsal-root-ganglion (DRG) section & Thy1-GFP-positive mouse brain section, respectively, with FOV of 1×1 mm^2^, scale bar of 150 µm; (b) & (d) noise-suppressed contrast-enhanced results with (a) α_max_ = 8.0 & (c) α_max_ = 6.0, respectively; R1-3 and R4-6: 458×458-sized and 490×490-sized regions-of-interest (ROIs), respectively, cropped from (a) & (b), and (c) & (d), respectively, with scale bar of 15 µm; C1-6: 50×50-sized ROIs cropped from red-arrow marked locations in R1-6; (e) intensity profiles along L1-6 (in C1-6), and (f)-(h) SNR, SBR & contrast-ratio plots for R1-6; (e)-(h) validate significant improvements on SNR, SBR and contrast-ratio, red & gray colors indicate before-& after-process cases, respectively.

To visualize the effectiveness of our approach, an adequate magnification is required. We thus perform a 12× digital zoom, and crop out three 458×458-sized regions-of-interest (ROIs) from the original 5500×5500-pixeled Nav1.8-tdTomato-image, marked as R1-3 in Figures 2(a) and (b) with unique colored-dashed-boxes. The magnified ROIs are shown on the right side of Figure 2(b), each with a scale bar of 15 µm. Likewise, another three 490×490-sized ROIs from the original 5926×6000-pixeled Thy1-GFP-image are marked as R4-6 in Figures 2(c) and (d), which are zoomed alongside, each with a 15 µm scale bar. At this point, our method’s effectiveness can be visualized with an observation of the two ROI-columns for R1-6, indicating before- and after-processing scenarios. Remarkably, the cell bodies in R1-3 are enhanced, yet well-preserved against saturation while enhancing the nearby weaker fibers.

To better study the effect, we select 50×50-sized ROIs from the red-arrow-marked locations in the before- and after-process sets of ROIs (R1-6), which are again zoomed as C1-6 sequentially. In Figure 2(e), we now plot intensity profiles along the red-dotted lines L1-6 (marked in C1-6), where red and gray colors indicate before (INP) and after-process (OUT) cases. We observe that in each case (L1-6), our approach effectively suppresses the noise contamination and drastically improves the contrast of the fine and weak neuronal structures.

Further extending our demonstration, two 5×5-sized ROI-1 and ROI-2 are taken from a signal location and a background location, respectively for each case of R1-6. For each ROI-1, we calculate the mean (μ_ROI−1_), and for each ROI-2, we calculate the mean (μ_ROI−2_) and standard deviation (σ_ROI−2_), and define SNR, SBR, and contrast-ratio as μ_ROI−1_ / σ_ROI−2_, μ_ROI−1_ / μ_ROI−2_, and ((μ_ROI−1_ −μ_ROI−2_) / (μ_ROI−1_ +μ_ROI−2_))×100%, respectively. Following these definitions, the SNRs, SBRs, and contrast-ratios for R1-6 are evaluated and plotted in Figures 2(f)-(h), respectively. The red and gray bars stand for before (INP) and after-process (OUT) scenarios, respectively. These plots essentially validate significant improvements on SNRs, SBRs and contrast-ratios for all the cases. Please note that, for a consistent analysis, we will be using the same ROI- and line-locations, and the same definitions of SNR, SBR, and contrast-ratio throughout the following analysis of our paper

### Demonstration of single-parametric control: effect of *α*_*max*_ over signal-to-noise ratio (SNR), signal-to-background ratio (SBR), and contrast-ratio

To quantitatively visualize the effect of *α*_*max*_ over SNR, SBR and contrast-ratio, we demonstrate Figure 3. From Figure 2, we take the same ROIs R1 and R2 for the Nav1.8-tdTomato image, and R4 and R5 for the Thy1-GFP image. The first row (INP) in Figure 3(a) shows the unprocessed ROIs R1-2 and R4-5, sequentially. Now, we gradually increase *α*_*max*_ from 3.0 to 8.0, and corresponding outputs are depicted in succeeding rows. The same sets of 5×5-sized ROIs as stated in Figure 2 are considered and corresponding values of SNR, SBR, and contrast-ratio are evaluated for R1-2 and R4-5 for each case of unprocessed input (INP) and corresponding outputs for *α*_*max*_ values of 3.0, 4.0, 5.0, 6.0, 7.0, and 8.0.

**Figure 3.**
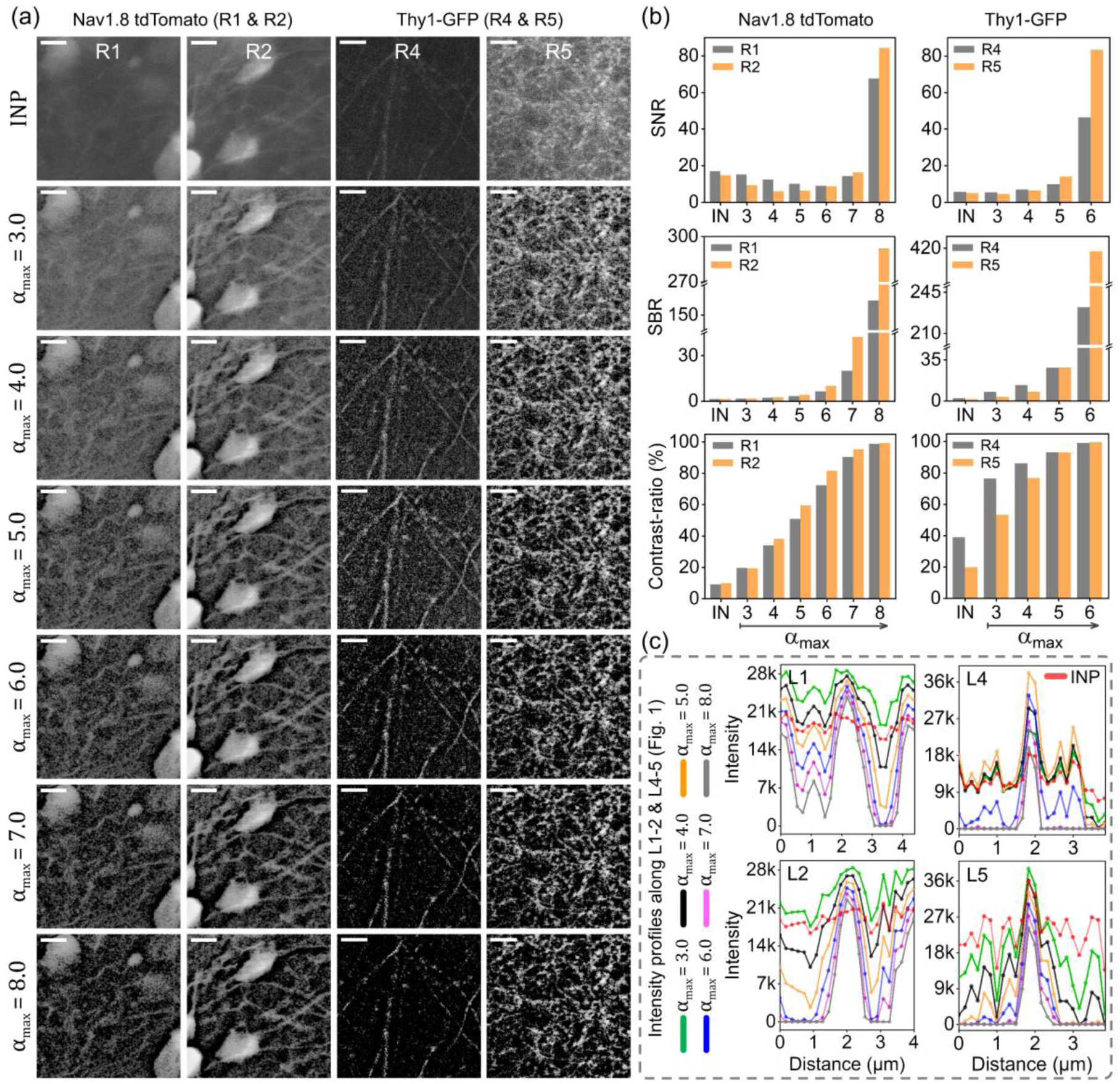
Demonstration of single-parametric control: effect of α_max_ over SNR, SBR & contrast-ratio. (a) ROIs R1-2 & R4-5 (from Figure 2), for unprocessed case (INP), and processed cases with α_max_ values of 3.0, 4.0, 5.0, 6.0, 7.0 & 8.0, respectively, scale bar = 15 µm; (b) SNR, SBR & contrast-ratio plots for R1-2 & R4-5 in first & second columns, respectively, SNR & SBR show exponential improvement as α_max_ goes above 6.0 in R1-2, and 4.0 in R4-5, contrast-ratio improves gradually for α_max_ ≥ 3 in each case; (c) intensity profiles along L1-2 & L4-5, plotted for INP (red), and respective outputs with α_max_ values of 3.0 (green), 4.0 (black), 5.0 (orange), 6.0 (blue), 7.0 (magenta) & 8.0 (gray), demonstrating simultaneous noise-suppression and contrast enhancement with increased α_max_.

In Figure 3(b), we plot the SNRs, SBRs, and contrast-ratios for R1-2 and R4-5 in the first and second columns, respectively. We observe that as *α*_*max*_ gradually increases, all three parameters SNR, SBR, and contrast-ratio tend to improve. SNR and SBR show an exponential improvement as *α*_*max*_ goes above 6.0 in case of R1-2, and 4.0 in case of R4-5. The contrast-ratio tends to improve gradually for *α*_*max*_ ≥ 3, for each case. We do observe that both σ_ROI−2_ and μ_ROI−2_ as defined in II.B become zero for *α*_*max*_ ≥ 9.0 in case of R1-2, and *α*_*max*_ ≥ 7.0 in case of R4-5. Continuing our assessment, Figure 3(c) plots the intensity profiles along L1-2 and L4-5 (see Figure 2, C1-2 and C4-5) for each case of INP (red) and respective outputs with *α*_*max*_ values of 3.0 (green), 4.0 (black), 5.0 (orange), 6.0 (blue), 7.0 (magenta), and 8.0 (gray). These intensity profiles essentially illustrate the progress of simultaneous noise-suppression and contrast enhancement with increasing *α*_*max*_. A simple observation of the red and gray curves in L1-2, and red and blue curves in L4-5 justifies the effectiveness of our method.

### Comparison with a few alternative software-based enhancement techniques

To ensure a fair comparison, we apply several alternative image processing methods to the full-FOV uncropped images of Nav1.8-tdTomato and Thy1-GFP samples previously shown in Figures 2(a) and (c), respectively. For a convenient visualization however, we consider the same ROIs R1, R2, and R3 (marked in Figure 2) and crop them out. The first row (1) in Figure 4(a) shows these three ROIs, each with a scale bar of 15 µm. The subsequent rows in Figure 4(a) show the results of (2) multiplicative gain enhancement, (3) minimum-maximum range adjustment, (4) histogram equalization (HE), (5) contrast-limited adaptive histogram equalization (CLAHE), (6) unsharp masking (UM), (7) morphological erosion, (8) morphological opening, (9-10) rolling-ball and sliding-paraboloid background subtractions, and finally (11) our proposed method, for each case of R1-3. Now, following the same sets of 5×5-sized ROIs used in Figure 2, we evaluate the SNR, SBR, and contrast-ratio for each case/ method (1-11) and for each ROI (R1-3), and subsequently plot them in Figure 4(b), where gray, orange, and cyan bars denote the results for R1, R2, and R3, respectively. Further extending our comparison, in Figure 4(c), we plot the intensity profiles along L1-3 (see Figure 2, C1-3) for each case of (1-11).

**Figure 4.**
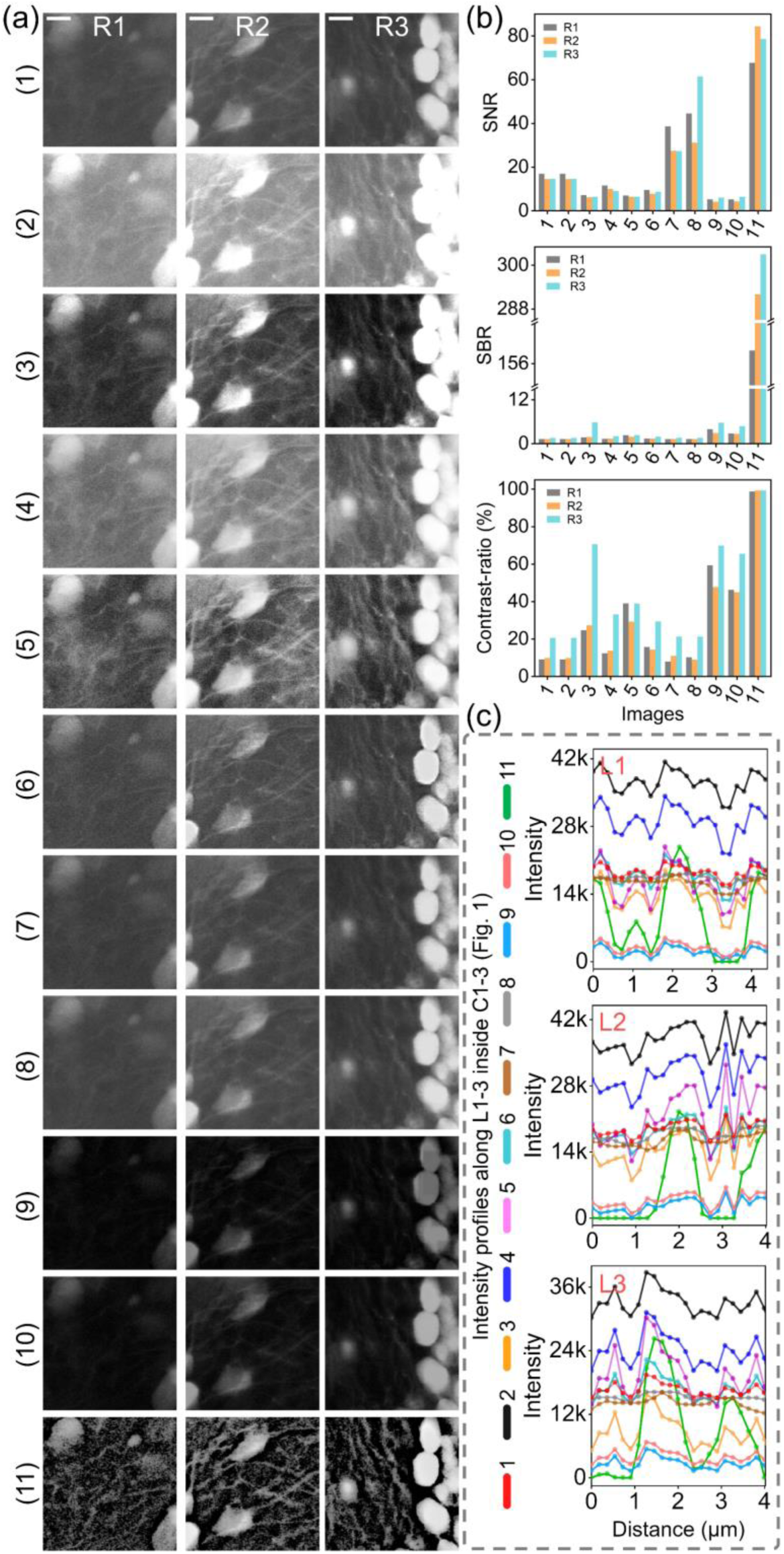
Comparison with a few alternative software-based enhancement techniques. (a) ROIs R1, R2 & R3 (from Figure 2) with scale bar of 15 µm, depicted for (1) INP (unprocessed), (2) multiplicative gain enhancement, (3) minimum-maximum range adjustment, (4) histogram equalization (HE), (5) contrast-limited adaptive histogram equalization (CLAHE), (6) unsharp masking (UM), (7) morphological erosion, (8) morphological opening, (9-10) rolling-ball and sliding-paraboloid background subtractions & (11) our proposed method; (b) SNR, SBR & contrast-ratio plots for (1-11), gray, orange & cyan bars denote results for R1, R2 & R3, respectively; (c) intensity profiles along L1-3 plotted for (1-11). Contradicting (2-10), our method (11) enhances SNRs, SBRs & contrast-ratios all at the same time. While enhancing weaker structures, our method prevents saturation to the brighter ones, for instance, cell-bodies in R3 are mostly saturated in (2) & (3), whereas they are well preserved in (11).

From both analysis in Figures 4(b) and (c), we observe that for morphological erosion (7) and opening (8), the SNRs tend to increase as they reduce high-frequency noise information resulting in a lower noise standard deviation, however, SBRs and contrast-ratios do not show a substantial improvement. In case of rolling-ball and sliding paraboloid background subtractions (9-10), the contrast-ratios tend to improve, however, the signal information tend to reduce simultaneously in each case (Figure 4(c), 9-10), and no substantial SNR- and SBR-improvements are observed. Likewise, CLAHE (5) improves the contrast-ratios, however, often encounters noise-amplification and thus results in poor SNR and SBR. Contradicting such approaches (2-10), our approach (11) successfully enhances SNRs, SBRs, and contrast-ratios all at the same time. It is further remarkable that while enhancing the visibility of weaker structures, our approach prevents saturation to the brighter ones. For instance, the bright cell-bodies in R3 have mostly been saturated in case of (2) and (3), whereas they are well preserved in our case (11).

### Assessment and comparison of time-complexity, and validation of real-time applicability

Figure 5(a) plots the average processing time for our proposed method in milliseconds with respect to the input image-size (16-bit). The red curve depicts average processing time via a conventional CPU i7-9800X consuming up-to ∼2000 ms for a 10000×10000-sized 16-bit input. The blue and green curves indicate average processing time via two CUDA-enabled GPUs, Quadro P1000 and Quadro RTX 8000, with CUDA-core numbers of 640 and 4608, respectively. Both these GPUs show a significant improvement to the processing speed. For instance, RTX8000 consumes only ∼111 ms for a 10000×10000-sized 16-bit input, which seems to be around 18 times performance boost in comparison to 9800X. Likewise, for a 1000×1000-sized 16-bit input, processing time for 9800X is ∼21 ms, whereas RTX8000 takes <3 ms for the same.

**Figure 5.**
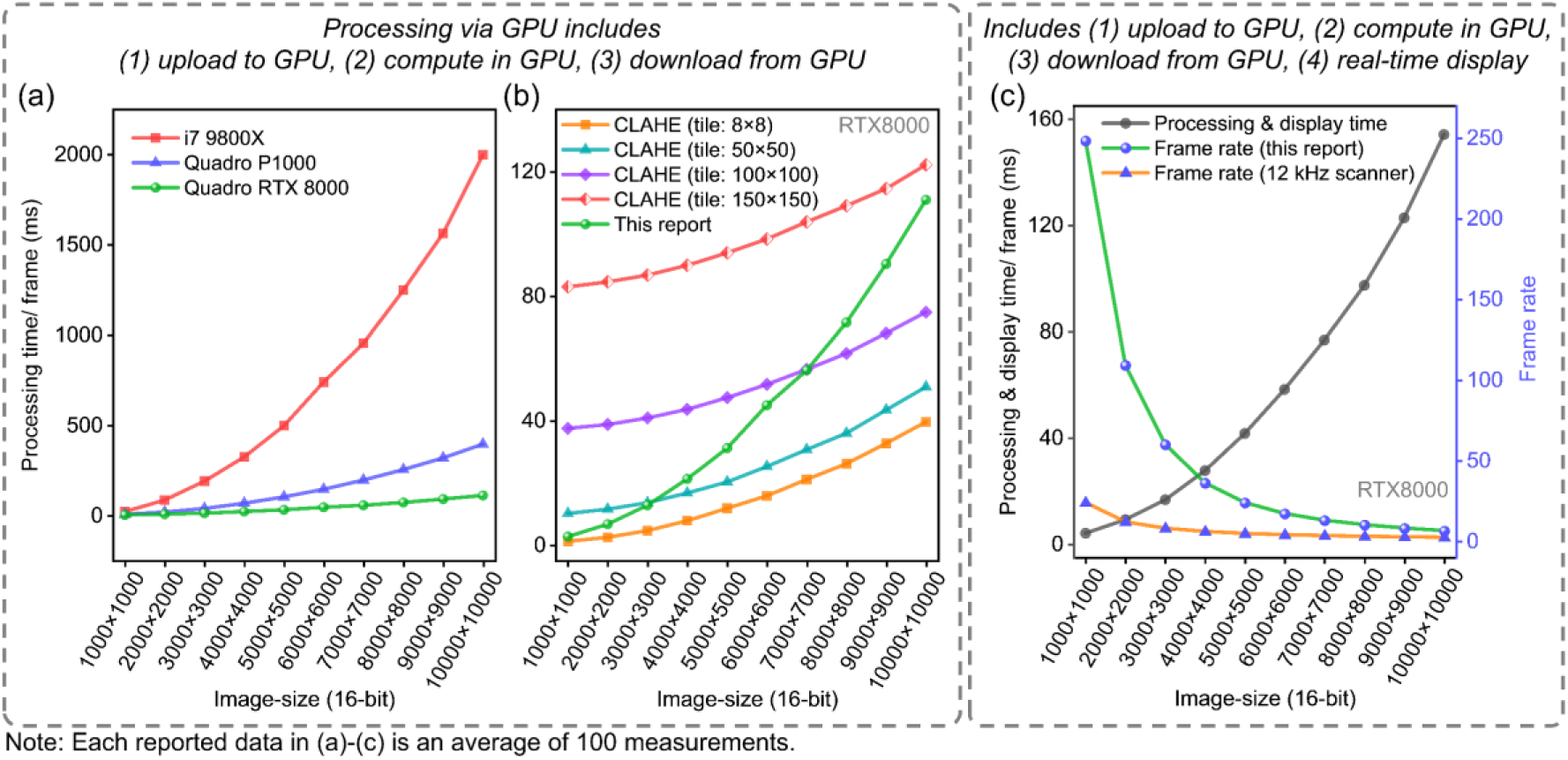
Assessment and comparison of processing speed, and validation of real-time applicability. (a) Average processing time in milliseconds per frame plotted for i7-9800X (red), Quadro P1000 (blue) & Quadro RTX 8000 (green), with respect to input image-size (16-bit); for 10000×10000-sized 16-bit input, 9800X consumes ∼2000 ms, whereas RTX8000 takes ∼111 ms with an 18× performance boost. (b) Average processing time in milliseconds per frame with respect to 16-bit input size, plotted for contrast limited adaptive histogram equalization (CLAHE) at different tile-grid-sizes & our proposed algorithm; CLAHE performance seems to reduce for larger tile-grid-size, whereas proposed algorithm is consistent for any α_max_. (c) Average of total processing & display time in milliseconds per frame (left axis), and respective frame-rate (right axis) for our proposed algorithm plotted with respect to 16-bit input size; observed frame rate (green) is substantially higher than maximum mechanical frame-rate for a state-of-the-art 12 kHz resonant scanner (orange).

We now compare our processing time with a widely used state-of-the-art technique, contrast limited adaptive histogram equalization (CLAHE). For a reasonable comparison, both CLAHE and our proposed method are tested through RTX 8000 with 16-bit input images. In Figure 5(b), the green curve plots the average processing time for our algorithm, while the others plot the same for CLAHE at different tile-grid-sizes. It is observed that for smaller tile-grid-size, CLAHE seems to be faster than our proposed method, however, its performance reduces gradually as we increase the tile-grid-size. On the other hand, for a particular input size, our algorithm’s performance is consistent for any recommended value of *α*_*max*_. Please note that each reported GPU-processing time in Figures 5(a) and (b) includes uploading the input data from host to GPU, processing the data in GPU, and downloading the output data from GPU to host.

Up-to this stage, we do not consider displaying the downloaded (from GPU) data to a computer screen. To assess the effective performance of our method, we now implement a real-time displaying of the processed images to a monitor, and measure the total time required for uploading to GPU, processing in GPU, downloading from GPU, and displaying to a monitor, for different input image sizes. The black and green curves in Figure 5(c) plot the total time required per frame and the corresponding frame rate, respectively, while varying the input data sizes. We observe that for a 10000×10000-sized 16-bit input data-set, a total time of ∼154 ms is consumed, thus resulting in a frame rate of >6 frames per second (fps). Likewise, for a 1000×1000-sized 16-bit input, the total time per frame is found to be ∼4 ms, and thereby a frame rate of ∼248 fps becomes feasible. Please note that for our analysis, we employed different image sizes from 1000×1000 up-to 10000×10000. However, our display screen was limited with a resolution of 3840×2160 and a refresh rate of 60 Hz.

For a high-speed laser-scanning fluorescence microscopy, researchers often employ fast enough resonant scanners to facilitate a high-speed raster scanning. If we now consider a state-of-the-art ultra-fast resonant scanner with a 12 kHz scanning frequency (24 kHz line rate), the maximum achievable mechanical frame rate with a bi-directional scanning is represented by the orange-curve in Figure 5(c), where we observe a frame rate of 2.4 fps and 24 fps for slow-axis line numbers of 10000 and 1000, respectively. It is thus evident that our effective frame rate is remarkably higher than the maximum allowed mechanical frame rate for such a 12 kHz scanning system, allowing our approach to be implemented in real-time imaging applications without any frame-lapse.

## Discussion

In this paper, we report a dedicated-hardware-free rapid noise-suppressed wide-dynamic-range compression and/or contrast enhancement method to virtually mimic the effect of a hardware-based adaptive illumination technique. Performing large-FOV aliasing-free two-photon fluorescence neuronal imaging at multiple excitation wavelengths, we validate the effectiveness of the proposed method by retrieving weak-intensity ultrafine neuronal structures amidst a strong noisy background, while demonstrating simultaneous improvements to the SNR, SBR, and contrast-ratio. A CUDA-assisted reduced time-complexity of <3 ms for a 1000×1000-sized 16-bit data-set enables a real-time applicability of the same.

For a better performance, the input image is expected to be not saturated. One can adjust excitation power, gain of the fluorescence detection system as well as input-range of the digitizer in order to prevent saturation. The proposed method tends to suppress a noisy background near the edges of a bright structure more aggressively compared to that near a weaker structure. That is to say, the contrast of the bright structures will be boosted first even at a lower *α*_*max*_, and as the value of *α*_*max*_ is increased, the contrast of the weak structures will gradually improve. For a lower value of *α*_*max*_, this behavior therefore might lead to an artifact particularly in case of a strong noisy background coexisting with bright enough or saturated structures. To minimize the same, a higher value of *α*_*max*_ is necessary. However, an excessively higher *α*_*max*_ might tend to suppress useful low-frequency information along with the noisy background. Based on our observations, we recommend an *α*_*max*_ range of 3.0 to 8.0. Practically, one should first apply a lower *α*_*max*_ value, and based on visual perception, *α*_*max*_ should be increased until a satisfactory result is observed.

We recommend that the data-acquisition system satisfies or exceeds the respective Nyquist-Shannon criterion [61-62], so that a smallest resolvable structure gets digitized with at least 4 pixels. One should not downscale the data prior to applying our method, however, if the sampling pixel-size is much smaller than that required by the Nyquist-Shannon criterion, a suitable pixel-binning can be performed. It is recommended not to apply a conventional low-pass filter to the digitized data-set, prior to applying our method, as the noise components will tend to lose their high-frequency nature and our approach might treat them as low-frequency information thereafter. Our approach boosts the weaker structures while preventing the brighter ones from getting saturated. Such local enhancement however might not be suitable to be applied in certain quantitative analysis.

The reported method can be utilized in a conventional or mesoscopic optical microscopy system to enhance the overall contrast, SNR, and SBR, and thereby to improve the image fidelity. Especially for a large-FOV imaging scenario, a wide variation in structural texture and signal strength is very likely to occur, where our proposed method can perform a significant improvement. For an extended-FOV imaging, our method can help to improve the visibility of FOV-edges and corners, which can further facilitate an artifact-free digital-image-stitching while imaging an ultra-large centimeter-scale tissue-sample. For a volumetric imaging application, the proposed method can significantly improve the visibility of finer and weaker structures particularly at a high enough penetration depth. The reported method has a tremendous potential to be applied prior to segmentation of neuronal structures, which can help construct ultra-high-resolution 3D-neuronal-maps of ultra-large volumetric brain regions, or even an intact whole animal brain.

In this paper we have demonstrated our idea by means of two-photon fluorescence images only, however, the same can be further extended to other forms of optical linear and nonlinear imaging, as well as clinical applications, such as, ultrasound, computed tomography, X-ray, and magnetic resonance imaging.

## Acknowledgments

This project was supported by Ministry of Science and Technology (Taiwan) with financial grants MOST 110-2321-B-002-011. We thank Dr. Daniel Lin (SunJin Lab Co., Taiwan), Dr. Jye-Chang Lee, and Prof. Chen-Tung Yen (Department of life science, National Taiwan University, Taipei 10617, Taiwan) for providing us with biological *ex vivo* samples for imaging.

## Author Contributions

B. J. Borah developed the method, wrote C++ codes, performed experiments and data analysis. C.-K. Sun supervised the project. B. J. Borah and C.-K. Sun wrote the paper.

## Method details

### Biological samples

The animals used in this study for preparation of the *ex vivo* test-samples were maintained in accordance with guidelines approved in the Codes for Experimental Use of Animals of the Council of Agriculture of Taiwan, based on the Animal Protection Law of Taiwan. All experimental protocols were approved by the Institutional Animal Care and Use Committee of National Taiwan University, Taipei, Taiwan.

### Two-photon fluorescence imaging of neuronal structures

Two-photon fluorescence imaging [63-64] was performed using a high-NA (>0.9) and low magnification (20×) objective lens (Olympus XLUMPlanFl, 20×/0.95W). The scanning head employed a resonant scanner (CRS 4 kHz, driver: 311-149887, Cambridge Technology, MA, USA) and a galvanometer scanner (8320K, driver: MicroMax 671, Cambridge Technology, MA, USA) for fast- and slow-axis scanning, respectively. A custom tube lens and a general scan lens (LSM05-BB, Thorlabs, NJ, USA) with effective focal lengths of 167 mm and 110 mm, respectively, were used providing a ∼1.5× beam magnification.

A 70 MHz, <60 fs fiber laser (Fidelity-2 Fiber Laser, Coherent, Inc., CA, USA) centered at 1070 nm was utilized directly with a one-pulse-per-voxel acquisition scheme for excitation of the Nav1.8-tdTomato sample. For excitation of the Thy1-GFP sample, the central wavelength was shifted to ∼919 nm. To achieve the same, output from Fidelity-2 fiber laser was free-space-coupled to a 7 mm-long photonic crystal fiber to induce a negative dispersion to generate Cherenkov Radiation [65-66]. Long-pass and short-pass filters with cut-on and cut-off at 750 nm and 1000 nm, respectively, were used to ensure a spectrum centered around 919 nm. A pulse duration of <60 fs was ensured after the objective lens by means of pulse pre-chirping [67] with a grating-pair.

A dichroic beam splitter (FF735-Di02, Semrock) was used to reflect the emerging fluorescence signal into a detection unit comprising a relay system with two lenses with effective focal lengths of 150 mm (Edmund Optics: 32-982) and 40 mm (Edmund Optics: 48-654), respectively, producing a 3.75× demagnification, and subsequently a photomultiplier tube (PMT, R10699, Hamamatsu Photonics, Japan). To ensure detection of Nav1.8-tdTomato and Thy-1 GFP two-photon fluorescence signal, two band pass filters, FF01-580/60-25-D, and FF03-525/50-25, respectively, were placed before photo-sensitive area of the PMT. A colored glass filter (FGB37-A, Thorlabs) was additionally placed in series with the band-pass filter in each case. A transimpedance amplifier C6438-01 (Hamamatsu Photonics, Japan) was employed to perform current to voltage conversion of the PMT-output signal, which was subsequently digitized with a high-speed digitizer ATS9440 (Alazar Technologies Inc., Canada).

### Implementation of the method via GPU-acceleration

For GPU-accelerated image/data processing, an open-source computer vision library, OpenCV (version: 4.5.0) was built with CUDA libraries (version: 10.1, update 2).

## Data processing

All intensity profiles along L1-6 shown in Figures 2, 3, and 4 were obtained via ImageJ (1.53c) software. In Figure 4, a multiplicative gain enhancement of 2.0 was used in (2); minimum to maximum range was set as 20% to 60% of the maximum (i.e., 65535) in (3); a tile-grid-size of 20×20 and a clip-limit of 4.0 were used for contrast-limited adaptive histogram equalization (CLAHE) in (5); a radius and a mask-weight of 7-pixels and 0.5, respectively, were used for unsharp masking (UM) in (6); a 5-pixel wide elliptical structure element was employed for both morphological erosion and opening operations in (7) and (8), respectively; a ball-radius of 70-pixels was used for rolling-ball and sliding-paraboloid background subtractions (with enabled smoothing) in (9) and (10), respectively. Each reported processing time in Figures 5(a)-(c) is an average of 100 measurements performed with standard C++ functions. CUDA (10.1)-accelerated OpenCV (4.5) built-in functions were employed for both methods in Figures 5(b). In each GPU-processing case in Figures 5(a)-(c), asynchronous data-transfer was employed in between host and GPU.

